# Significance Analysis for Clustering with Single-Cell RNA-Sequencing Data

**DOI:** 10.1101/2022.08.01.502383

**Authors:** Isabella N. Grabski, Kelly Street, Rafael A. Irizarry

## Abstract

Unsupervised clustering of single-cell RNA-sequencing data enables the identification and discovery of distinct cell populations. However, the most widely used clustering algorithms are heuristic and do not formally account for statistical uncertainty. Many popular pipelines use *clustering stability* methods to assess the algorithms’ output and decide on the number of clusters. However, we find that by not addressing known sources of variability in a statistically rigorous manner, these analyses lead to overconfidence in the discovery of novel cell-types. We extend a previous method for Gaussian data, Significance of Hierarchical Clustering (SHC), to propose a model-based hypothesis testing approach that incorporates significance analysis into the clustering algorithm and permits statistical evaluation of clusters as distinct cell populations. We also adapt this approach to permit statistical assessment on the clusters reported by any algorithm. We benchmarked our approach on real-world datasets against popular clustering workflows, demonstrating improved performance. To show its practical utility, we applied it to the Human Lung Cell Atlas and an atlas of the mouse cerebellar cortex. We identified several cases of over-clustering, leading to false discoveries, as well as under-clustering, resulting in the failure to identify new subpopulations that our method was able to detect.

## 1 Introduction

Unsupervised clustering is widely applied in single-cell RNA-sequencing (scRNA-seq) workflows. The goal is to detect distinct cell populations that can be annotated as known cell-types or discovered as novel ones. A common claim from applying these workflows is that new cell subtypes or states have been discovered because a clustering algorithm divided a population associated with a known cell type into more than one group. But how do we know if this partition could have happened by chance, even with only one cell population being present? Current approaches do not consider this question. Furthermore, because the most popular clustering algorithms are heuristic and do not rely on underlying generative models, they are simply not designed for statistical inference.

As an example, consider the Louvain and Leiden algorithms [1] as implemented by the widely used Seurat toolkit [2]. A standard procedure is to 1) apply principal component analysis to the log-transformed and normalized counts, 2) compute the Euclidean distance between the first 30 principal components of each pair of cells, 3) find the 20 nearest neighbors for each cell, 4) specify a weight for each pair of cells based on the number of neighbors in common and use this to define weighted edges in a network, and 5) divide the network into clusters that maximize *modularity* [1].

The number of clusters found is related to a tuning parameter called the *resolution*, the best value of which is typically selected by manual inspection of the clusters or by maximizing criteria related to *clustering stability* [3, 4, 5, 6]. Note that an underlying generative model is not provided to motivate any of these steps nor to assess how much the results can change due to natural uninteresting random variation. Hence, these and other similar algorithms do not examine the statistical possibility of under- or over-clustering, which leads to the failure to detect rare populations or the false discovery of novel populations, respectively.

Over-clustering can be particularly insidious because clustering algorithms will partition data even in cases where there is only uninteresting random variation present [7]. Moreover, due to the data snooping bias, also known as double-dipping, cells that have been incorrectly clustered into two groups can have genes that appear to be differentially expressed with spuriously small *p*-values [8]. This is because, for example, if we force a single population into two clusters, the algorithm will assign cells that are more similar to each other to the same group, but the statistical test does not take this selection into account when considering the null hypothesis. As a result, if one does not account for this statistical reality, over-clustered output can appear to show convincing differences.

Statistical inference frameworks for clustering have been introduced in contexts other than cell population discovery with scRNA-seq data [9, 10]. These assume an underlying parametric distribution for the data, specifically Gaussian distributions, where distinct populations have different centers. A given set of clusters can then be assessed in a formal and statistically rigorous manner by asking whether or not these clusters could have plausibly arisen under data from a single Gaussian distribution. If so, then the set of clusters likely indicates over-clustering. However, a limitation of many of these approaches in the context of scRNA-seq cell population discovery is that one can only compare one versus two clusters, rather than any number of clusters, and clustering cannot be done in a hierarchical fashion. Significance of Hierarchical Clustering (SHC) [11] addresses this limitation by incorporating hypothesis testing within the hierarchical procedure. However, SHC is not directly applicable to scRNA-seq data due to the Gaussian distributional assumption, which is inappropriate for these sparse count data [12, 13].

Here, we extend the SHC approach to propose a model-based hypothesis testing framework embedded in hierarchical clustering for scRNA-seq data. Motivated by previous exploratory analyses [13], we defined a parametric distribution to represent cell populations, and developed an approach implemented in two ways. First, like SHC, our approach can perform hierarchical clustering with built-in hypothesis testing to automatically identify clusters representing distinct populations. We refer to this self-contained method as single-cell SHC (sc-SHC). To permit significance analysis for datasets that have already been clustered, we developed a version that can be applied to any provided set of clusters. Our approach corrects for multiple, sequential hypothesis testing and controls the family-wise error rate (FWER), with interpretable summaries of clustering uncertainty. We motivate the need for statistical inference in scRNA-seq clustering pipelines, describe the mathematical details of our approach, benchmark our approach on real data against popular clustering workflows, and finally demonstrate the advantages with both in the Human Lung Cell Atlas and a mouse cerebellum dataset.

## 2 Results

### 2.1 Current clustering workflows over-cluster

To assess the performance of the *clustering stability* approach applied in current workflows to avoid over-clustering, we simulated scRNA-seq data from a single distribution representing one cell population (Figure 1A). Because typical clustering workflows only consider a subset of high-variance genes, we simulated 5,000 cells with 1,000 genes each, intended to represent expressed genes, using a previously published model [13] (see Section 1). The expression counts for each cell were drawn from the same joint distribution over all 1,000 genes, and thus an algorithm that reports more than one cluster in these data is over-clustering.

**Figure 1:**
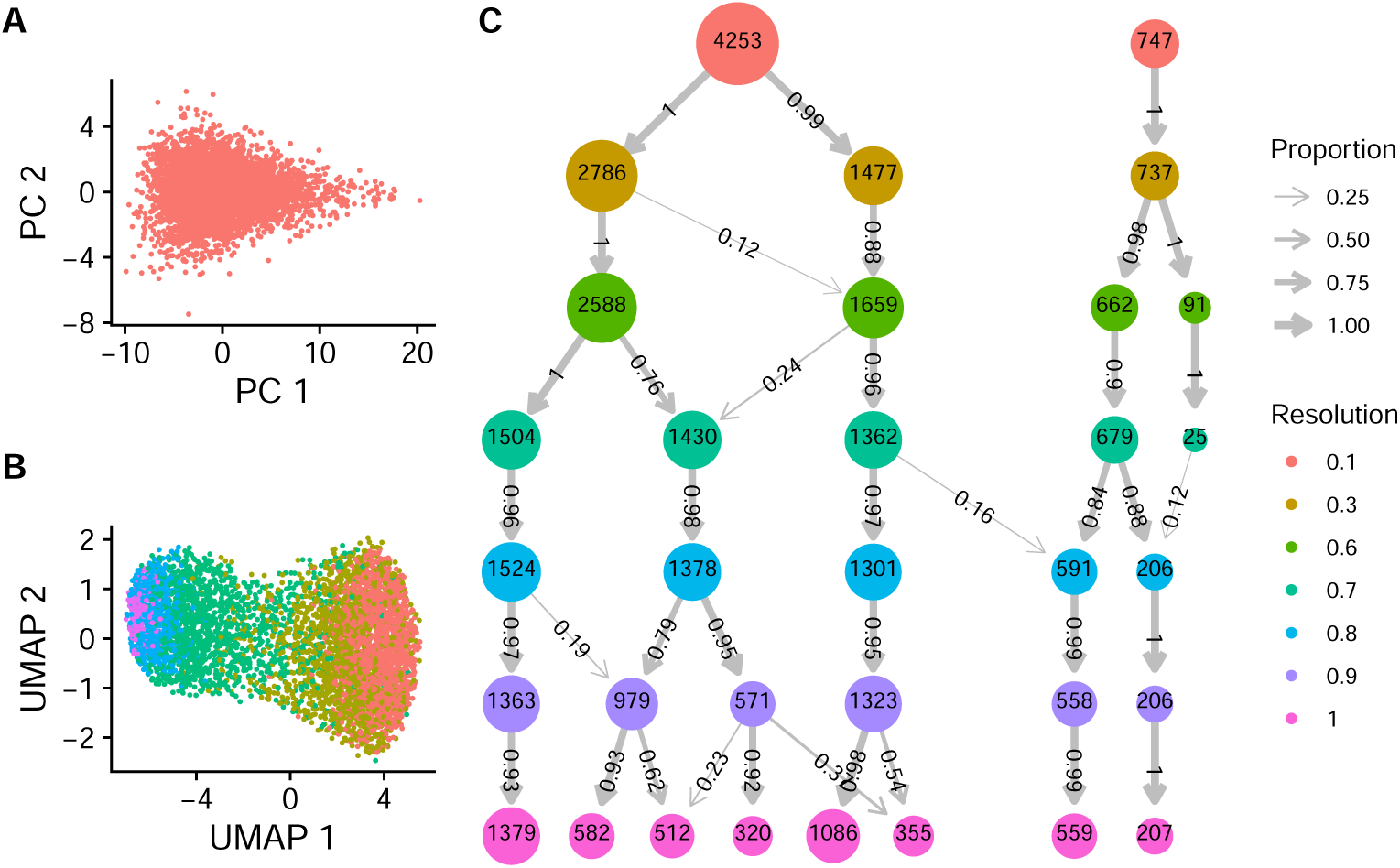
Clustering results for applying Seurat’s implementation of the Louvain algorithm to simulated data representing one cell population. (A) PCA plot of 5,000 cells generated from the same joint distribution over 1,000 genes. (B) UMAP plot colored by cluster membership when using the default resolution parameter of 0.8. (C) A modified `clustree` visualization of the resulting clusters using Seurat’s implementation of the Louvain algorithm at different resolution parameters, denoted by row. The number of circles in each row represents the number of clusters found with the respective resolution parameter, with the circle’s label and area denoting the number of cells in the cluster. The arrow widths and labels denote what proportion of the cells in each cluster came from the clusters at the previous resolution parameter.

We clustered these simulated expression counts using the most popular clustering algorithm, Seurat’s implementation of the Louvain algorithm [2] (see Supplementary Methods 1 for details). We compared results obtained using different *resolution parameters* used to control the final number of clusters. The number of clusters found varied from two to eight, with the default resolution parameter resulting in five clusters, demonstrating substantial over-clustering (Figure 1B). Furthermore, as the resolution parameter increased, most clusters represented increasing subdivisions of previous cluster output, indicating a high degree of stability (Figure 1C). This demonstrates that clustering stability does not necessarily provide a useful statistical assessment. Moreover, applying the standard approach of visualizing the clusters with UMAP plots provided further confirmation bias (Figure 1B). The current approach is therefore prone to over-clustering with no tools provided for detecting the problem.

### 2.2 Significance analysis can both identify and assess clusters

We introduce a model-based hypothesis testing framework that can be applied to any provided set of clusters to determine which, if any, should be merged due to over-clustering. The approach can also be built into hierarchical clustering [11] to produce a self-contained clustering pipeline that identifies clusters with statistical evidence that they represent distinct populations. The key idea is to define a realistic parametric distribution to represent a cell population, and then assess whether a separation into two clusters, proposed by the algorithm, could have arisen by chance from cells belonging to only one population. We use a parametric joint distribution that takes into account natural and technical variability, as well as correlation between genes (details in Section 4).

To illustrate the foundational concepts of our framework, we first describe the simplest case. Suppose we have a dataset that has been clustered into two groups, and we want to evaluate whether these two clusters should be just one. Specifically, we want to evaluate whether the clustering algorithm could have found such clustering structure, by chance, from data generated by a single distribution. To answer this question, we first computed a quality assessment metric for the cluster, specifically the average silhouette [14], which we denote with *s*. We then fit our parametric model, assuming just one population, to the data to obtain a *null* model. We used this model to perform a parametric bootstrap procedure (Figure 2A) [10] and form a null distribution for the average silhouette. With a null distribution in place, we can estimate a p-value: the probability of observing an average silhouette for two clusters as high as or higher than *s* when only one population is present.

**Figure 2:**
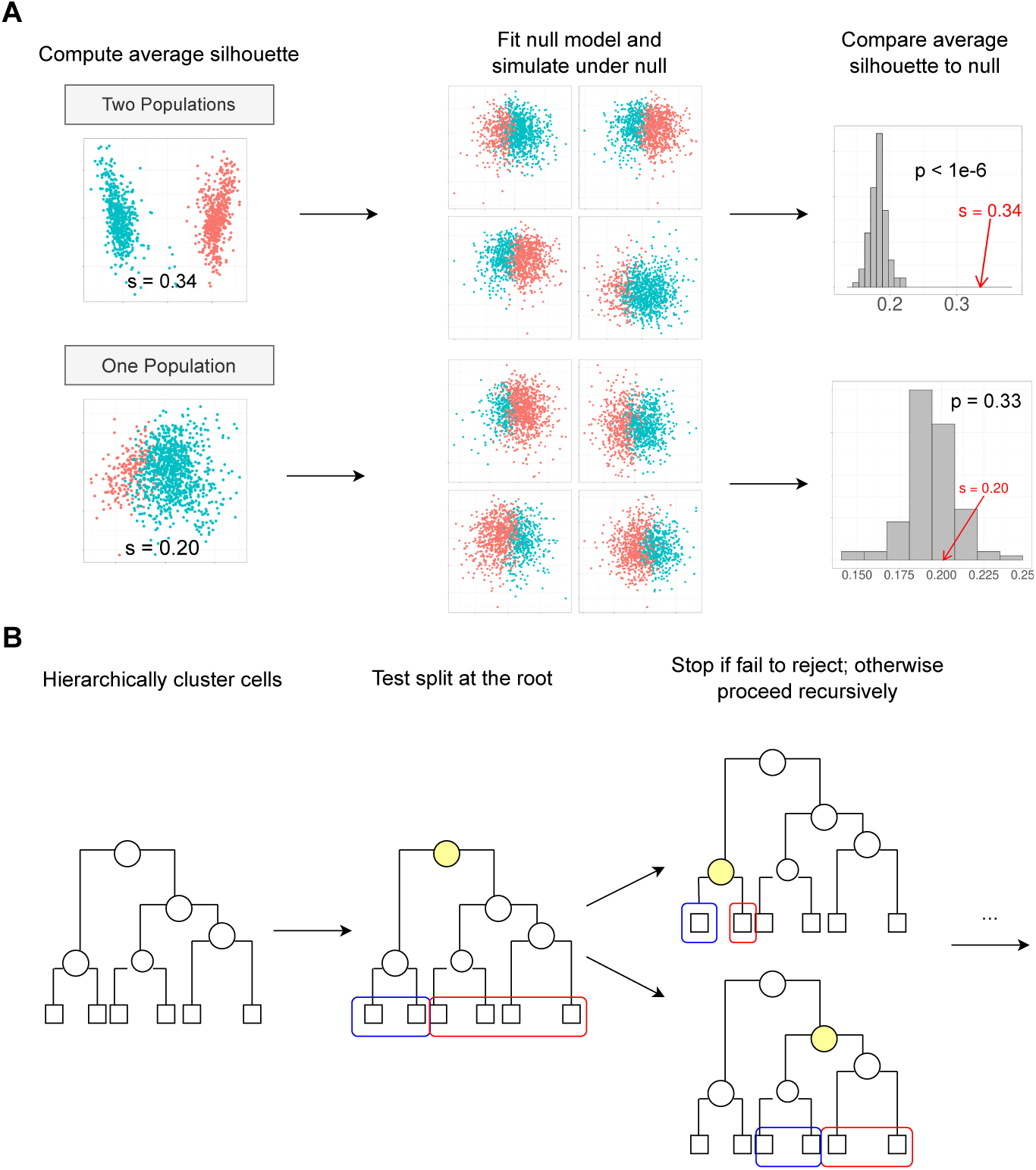
Schematic illustrating our approach to significance analysis for clustering. (A) Schematic of the test used by our approach to decide whether a proposed two-way split is significant. We show two examples, one in which two distinct populations were simulated (top) and one in which only one population was simulated (bottom). The schematics show how we use hierarchical clustering to divide the data into two, fit a single parametric model to the data, simulate 100 datasets under that model, and cluster each simulated dataset and compute the average silhouette. We then compare the average silhouette of the observed clusters to this empirical null distribution to decide whether to reject the null hypothesis. (B) Schematic of sc-SHC. We hierarchically cluster all the cells, and carry out significance analysis to decide whether to split the root node into the two clusters denoted by blue and red. We stop if we fail to reject the null hypothesis, and otherwise recursively continue performing tests to decide if we split each node or not.

To develop a clustering pipeline, we extended SHC [11] for scRNA-seq. Specifically, we applied hierarchical clustering to distances computed using a method developed for scRNA-seq (see Section 4). This resulted in a tree where the root node divided the cells into two groups, and each successive node encoded a further two-way split, all the way down to end nodes representing the individual cells. Note that we can obtain several different clustering results by merging branches [15]). To decide what branches to merge, we recursively applied the statistical test, described above, at each node, adjusting the significance threshold each time to control the family-wise error rate (FWER). After running this full procedure, we had a final set of clusters that have been determined by hypothesis testing to represent cells from distinct distributions (Figure 2B). We also provide a useful uncertainty summary for each cluster by adapting the concept of an adjusted p-value. Specifically, we run the pipeline using a FWER considered to be the highest tolerable false positive rate, then, for each split in the tree, we define an adjusted p-value as the infimum of FWER thresholds that would have permitted this particular split to be considered statistically significant. Full details are in Section 4.

To apply our significance analysis framework to any given set of pre-computed clusters produced by any algorithm, we modified our approach as follows. We first hierarchically clustered the centers of the provided clusters, such that each leaf of the tree represents one of the original clusters, and then again recursively applied our statistical test at each node. After running this procedure, we obtained a final, possibly merged set of clusters to correct for any potential over-clustering. Full details are in Section 4.

### 2.3 Significance analysis substantially improves current clustering approaches

To benchmark our method against current clustering workflows, we constructed three datasets with known ground truth: a *one-population* dataset, a *five-populations* dataset, and a *five-similar-populations* dataset. Specifically, we constructed our datasets using 2,885 cells from the 293T cell line [16]. Because these cells are all from the same cell line, they represent a single, well-defined population. The *one-population* dataset consists of these 2,885 cells, completely unaltered. The *five-populations* dataset consists of these same 2,885 cells, but modified to create five different populations of equal size, with up to 100 genes permuted to create differing gene expression levels across populations. Finally, the *five-similar-populations* dataset was modified in the same way except that the fourth and fifth populations differ only by ten genes, instead of 100 (see Supplementary Methods 5 for details).

We ran our clustering pipeline on each of these three datasets, as well as four popular existing clustering workflows for comparison: Seurat’s implementation of the Louvain algorithm (denoted here as *Seurat-Louvain*), Seurat’s implementation of the Leiden algorithm (denoted here as *Seurat-Leiden*) [2], Monocle’s `cluster_cells` function [17], and SC3’s consensus clustering algorithm [18]. Both Monocle and SC3 have built-in mechanisms to choose the number of clusters, but both Seurat-Louvain and Seurat-Leiden require the user to specify the resolution parameter, which is related to the number of clusters. We thus ran these latter algorithms in two ways: using the default resolution parameter, and using `clustree` to guide the choice of resolution parameter (see Supplementary Methods 5). We then also applied our significance analysis to the clustering outputs produced by each of these methods.

For each approach and dataset, we reported the number of clusters found and the Adjusted Rand Index (ARI), a metric of cluster accuracy where values closer to 1 indicate higher accuracy [19]. To compute the ARI, we compared the outputted cluster labels to the ground-truth population labels. In the one-population dataset, our clustering pipeline was the only method that correctly found just one cluster; other approaches found anywhere from two to fifteen clusters. While applying stability analysis (`clustree`) resulted in much improved performance compared to default parameters, these other algorithms still overestimated the number of clusters. However, applying our significance analysis improved the performance of all other algorithms (Table 1).

**Table 1.**
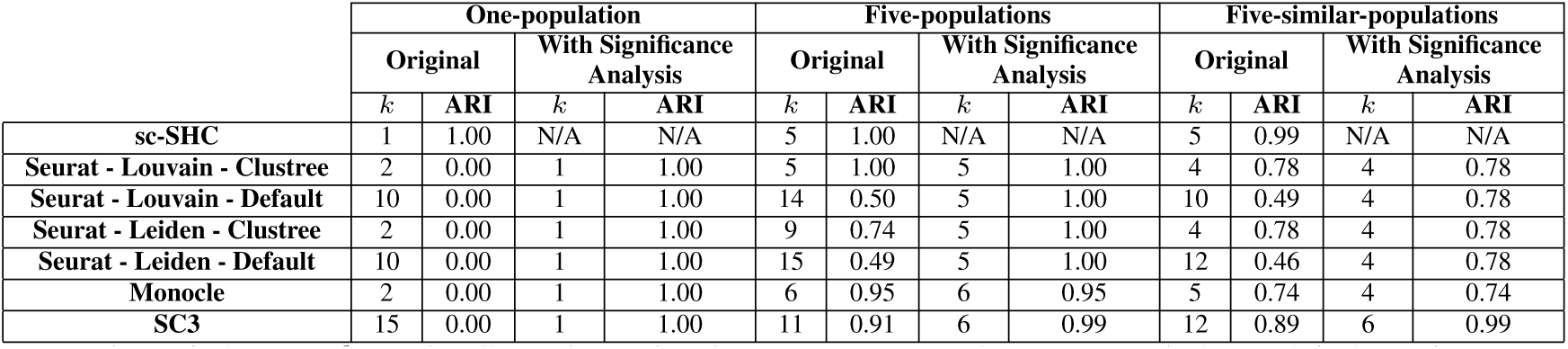
Number of clusters (*k*) and Adjusted Rand Index (ARI), comparing outputted cluster labels to the ground-truth population labels, when our clustering pipeline, sc-SHC, as well as existing methods, are applied to three datasets. We also applied our significance analysis on top of each existing method.

In the five-populations dataset, our approach again found the correct number of clusters (five) with perfect correspondence to the ground-truth labels. This was also achieved by Seurat-Louvain with `clustree`; all other approaches found between six and fifteen clusters. Applying significance analysis improved the performance of these methods. Specifically, the Seurat methods were merged to five clusters, again with perfect correspondence to the ground-truth labels. For SC3, significance analysis pruned the clusters from eleven to six and improved the ARI from 0.91 to 0.99. However, for Monocle, no clusters were merged (Table 1). We noted that the Monocle cluster outputs were not perfectly subdivided from the ground-truth labels, and further merges from the six-cluster solution would have mixed together cells from different populations, which is consistent with no improvements being possible given the starting clusters.

Finally, in the five-similar-populations dataset, our approach once again found the correct number of clusters (five), with an ARI of 0.99. Only one other approach, Monocle, found the correct number of clusters, but one of its clusters combined the two similar populations, resulting in a much lower ARI of 0.74. All other approaches either found too few clusters (four) or too many (between ten and twelve), but regardless of the number of clusters, all except SC3 combined the two similar populations into one. Applying significance analysis pruned SC3’s 12 cluster output down to six clusters with an ARI of 0.99. Although the improvement for the other methods was not as substantial, in all instances, applying significance analysis resulted in equal or better ARI (Table 1).

### 2.4 Significance analysis corrects over-clustering in the Human Lung Cell Atlas

To demonstrate how our approach provides different results in real-world data, we applied our significance analysis to clusters reported by the Human Lung Cell Atlas [20]. Specifically, we examined the 28,793 cells from the patient with the most cells available. The original study identified 39 clusters, some of which were interpreted as novel cell types. Our approach merged these clusters into 26, indicating over-clustering in the original project. Specifically, 17 of the 26 clusters found to be statistically significant corresponded exactly to 17 of the original clusters. The remaining 9 were combinations of two to four of the originally reported clusters.

To illustrate one of the examples of over-clustering, we consider what the original study reported as three clusters: Capillary Aerocytes, Capillary, and Capillary Intermediate 1. Applying significance analysis did not find evidence of the presence of the Capillary Intermediate 1 subpopulation and merged it with the Capillary cluster, while the Capillary Aerocytes were found to be a distinct population. An *scRNA-seq gating plot* (described in Section 4.4), comparing cells in the Capillary Aerocyte cluster to cells reported to be in the other two, shows strong evidence of two distinct modes, which was consistent with Capillary Aerocytes being a distinct population (Figure 3A). An *scRNA-seq gating plot* comparing the reported Capillary and Capillary Intermediate 1 clusters did not show evidence of two distinct populations (Figure 3B).

**Figure 3:**
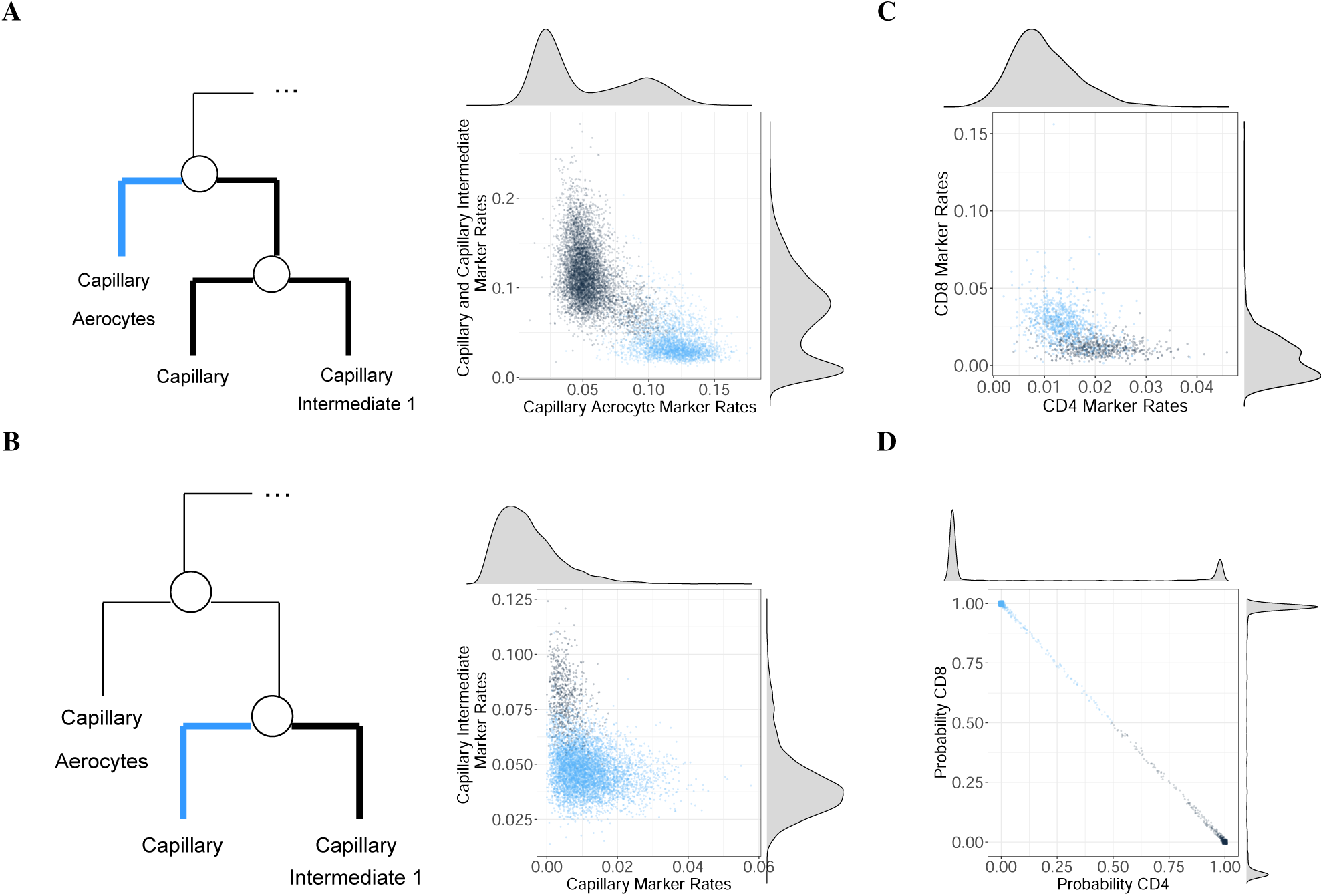
Applying significance analysis to clusters reported by the Human Lung Cell Atlas.(A) scRNA-seq gating plot for the proposed split between Capillary Aerocytes, and Capillary and Capillary Intermediate 1 cells. We compare the proportion of UMIs in each cell representing significantly more highly expressed genes in Capillary and Capillary Intermediate 1 cells to the proportion of UMIs representing significantly more highly expressed genes in Capillary Aerocytes. (B) scRNA-seq gating plot for the proposed split between Capillary and Capillary Intermediate 1 cells. (C) scRNA-seq gating plot comparing CD8 cells (comprising the CD8 Naive and CD8 Memory clusters, in blue) and CD4 cells (comprising the CD4 Naive and CD4 Memory clusters, in black). (D) Probability of each cell being assigned CD4 versus CD8 using a supervised annotation approach, with blue indicating new CD8 labels and black indicating new CD4 labels.

To illustrate further, we considered the originally reported CD4 Memory, CD4 Naive, CD8 Memory, and CD8 Naive clusters, which significance analysis merged into one cluster. Because these are widely studied immune cell types, we were able to independently validate the annotation using a reference-based cell annotation approach [13]. However, 18% of the cells were annotated differently by the clustering and reference-based approach. Furthermore, an scRNA-seq gating plot did not show obvious separation between the cells annotated as CD8 and CD4 by the clustering approach (Figure 3C). In contrast, the reference-based approach assigned probabilities to the cells in a way that formed two distinct modes (Figure 3D), indicating that distinct CD8 and CD4 populations were in fact present, but the clustering approach was unable to annotate them appropriately.

### 2.5 Our method identifies distinct subpopulations in the mouse cerebellum

We also applied our clustering pipeline to a mouse cerebellum dataset [21]. In particular, we examined 14,027 non-granule cells from the first female mouse cerebellar cortex, and found a total of 25 clusters. The original study clustered and annotated the data in an iterative process using both Seurat’s implementation of the Louvain algorithm [2] and LIGER [22]. They reported the resulting annotations at two levels of granularity – high-level major cell types, of which there are 17, and finer-level subclusters, of which there are 41. The ARI comparing our reported clusters with the major cell type level was 0.76, and the ARI with the subcluster level was 0.85, demonstrating that our clustering approach recovered overall similar structure as the original study.

We further examined some of the originally reported subclusters for which our significance analysis did not find supporting evidence. For example, the original study reports two astrocyte subclusters, whereas our approach found just one cluster containing all the cells originally labeled as astrocytes. This suggests that the original study’s finer-grained labels had over-clustered the astrocytes. The original study also reports nine Purkinje subclusters, of which two are specifically Purkinje anti-Aldoc subclusters and the remaining seven are Purkinje Aldoc subclusters. Our pipeline with significance analysis found only four clusters representing these Purkinje cells; one cluster contained 94% of cells from the original seven Purkinje Aldoc subclusters, while the other three clusters split up the Purkinje anti-Aldoc cells. This suggests that the original study had over-clustered the Purkinje Aldoc cells, but potentially did not subcluster the anti-Aldoc cells enough. We note that the adjusted p-value for splitting the Purkinje Aldoc cluster from the other

Purkinje cells was 4.6 *×* 10^*−*9^, indicating high certainty for separating Aldoc cells from anti-Aldoc cells. However, the adjusted p-value for the proposed split within the Purkinje Aldoc cluster, which did not achieve our significance threshold, was 0.35. The two splits that created the three Purkinje anti-Aldoc clusters had adjusted p-values of 0.056 and 0.075 respectively. Hence, at slightly more conservative FWERs, we would not have reported further substructure within the anti-Aldoc cells, which means there is considerable uncertainty around these additional clusters.

## 3 Discussion

Clustering analysis is an integral part of numerous scRNA-seq analyses pipelines. scRNA-seq data are affected by natural and technical random variability, yet the most popular pipelines do not account for statistical uncertainty. As a result, over-clustering is common, and overconfident interpretations can lead to flawed cell-type annotations and incorrect claims of discoveries of novel subtypes. To address this problem, we proposed a significance analysis framework that integrates algorithmic clustering with a probabilistic model. By assuming an underlying parametric model of gene expression, we built on previously developed statistical methodology to create a parametric bootstrap procedure that evaluates whether observed clustering structure can arise even when only one population is present. In particular, we presented two ways of applying this idea: a self-contained approach (sc-SHC) that builds hypothesis testing into hierarchical clustering to automatically identify clusters corresponding to distinct cell populations, and a *post-hoc* approach that can evaluate any provided set of clusters for possible over-clustering.

Using a simulation study, we demonstrated the substantial improvements provided by our approach when clustering with scRNA-seq data. In particular, we showed that current approaches are prone to over-clustering data and that performing significance analysis improves this by correctly merging spurious clusters. While stability analyses such as `clustree` yield notable improvement, they do not completely alleviate the problem of over-clustering. By contrast, in our simulations, sc-SHC found the correct number of clusters, with adjusted Rand indices above 0.99. We also showed that our approach prevented over-clustering without being overly conservative – even when two distinct populations differed only by the expression of ten genes, our approach still correctly separated them. Our *post-hoc* approach also performed favorably, reducing over-clustering in nearly every instance. However, we note that this post-hoc approach is limited by the accuracy of the original clustering: our approach can improve other methods only in reducing over-clustering, not in fixing incorrect annotations. Finally, to demonstrate that our approach makes a difference on real data, we applied this framework to the Human Lung Cell Atlas and a mouse cerebellum dataset to correct over-clustering and also identify previously missed distinct cell populations. Our approach also protected from overconfidence in a case in which two populations were present, and detectable with reference-based annotation tools, but the low signal to noise ratio made clustering unable to correctly form the two clusters. By merging these, our approach avoided the incorrect annotation of cells.

A limitation of model-based approaches is that the definition of a distinct population depends on the appropriateness of the parametric model assumed to describe gene expression. Although extensive data exploration indicates that the model we used does in fact describe observed data from distinct populations, scenarios in which it is not appropriate should be considered. For example, we assumed a unimodal marginal distribution for gene expression within a population. This implies that for cell populations in which some gene expression follows multi-modal distributions, our approach might also result in over-clustering as the incorrect assumption may lead to incorrectly rejecting the one population null hypothesis. Also note that, like all clustering algorithms, our approach identifies discrete populations of cells. Our model-based approach associates these discrete populations with different unimodal probability distributions. However, this does not preclude the presence of biologically meaningful variation within these populations. For example, a continuously varying gene expression level might be associated with different stages within a cell cycle. However, methods that partition cells into distinct groups, such as clustering, are not an appropriate tool for statistically describing these important sources of variability, although model-based approach such as ours can be adapted to quantify this variation through a continuous parameter in the model.

## 4 Methods

### 4.1 Significance Hierarchical Clustering for scRNA-seq

The first step in hierarchical clustering is computing a distance between each cell. Because scRNA-seq data are characterized by small counts and high dimensionality, computing Euclidean distance is not appropriate nor computationally convenient. We therefore compute Euclidean distance on the latent variables estimated by the approximate GLM-PCA procedure [12, 23] on the *G* = 2, 500 genes with the largest deviance under a multinomial null model [12]. We compute Euclidean distance on the first 30 latent variables to produce an *N × N* distance matrix *D*, with entries *d*_*i,j*_ representing the distance between cells *i* and *j* and *N* the number of cells. We then apply hierarchical clustering to *D* using Ward’s criterion [24].

To identify clusters corresponding to distinct populations from this tree, we followed a procedure similar to SHC [11]. We first decide on a desired family-wise error rate (FWER) of the entire procedure, referred to as *α*. For the results presented here, we used α = 0.05 in the simulations and *α* = 0.25 for the real data applications. We then proceeded to go down the tree, deciding which splits to keep and which to merge. We began at the root node, which can be interpreted as splitting all *N* cells into two clusters. We then decided whether or not to keep this split via hypothesis testing. First, we defined the average silhouette [14] as a *test statistic*, denoted as *s* and defined in detail in Supplementary Methods 4. Note that the better two clusters match the data, the higher the value of this statistic. Next, we estimated a null distribution for this statistic. To do this, we defined and fit a null parametric model ℱ, described in detail in Supplementary Methods 2, and then used parametric bootstrap to estimate a null distribution. Data exploration indicated that this null distribution can be approximated with a normal distribution, which permitted the use of fewer bootstrap samples to obtain a precise estimate of this distribution. With this estimate in place, we computed a p-value for *s*. Note that this assumption of normality can be avoided by obtaining more bootstrap samples and calculating the p-value from the resulting empirical distribution of bootstrapped *s* values.

If the p-value was greater than or equal to *α*, we failed to reject the null hypothesis, and concluded that all the data should belong to one cluster. Otherwise, we rejected the null hypothesis, split the data into the two proposed clusters, and continued recursively down the tree. Specifically, in the next step, we examined the next highest nodes, which propose further binary splits within each of these two newly formed clusters. We applied the same hypothesis test as above to each of these proposed splits. Because these proposed splits, unlike the split at the root node, pertain to subsets of the total cells, we made two modifications to the test. First, we re-applied the dimension reduction procedure to the relevant cells and computed an updated distance matrix for the average silhouette calculation. Second, to maintain the FWER at level α, we accounted for the multiple, nested nature of this hypothesis testing by comparing the p-value at a given node to 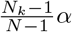, with *N*_*k*_ the number of cells below that node. This approach to FWER control was first proposed by [25] in the context of hierarchical variable selection, and was used by [11] in SHC.

When we failed to reject the null hypothesis at any given node, we did not test any further nodes on that branch. Instead, the cells below that node are all considered to belong to one cluster. Otherwise, if we did reject the null hypothesis, we continued testing at subsequent nodes. By the end of the procedure, we had a set of nodes where we failed to reject the null hypothesis, which corresponds to the final set of clusters.

### 4.2 Quantifying Uncertainty

We summarized the uncertainty for each split in the hierarchical cluster using the adjusted p-value metric. Specifically, at each split, we can report the infimum of FWER thresholds such that the split would have been found to be statistically significant. Suppose the p-value at node *k* is *p*_*k*_. To find the split statistically significant while controlling the FWER to level *α*, we require

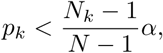

with *N*_*k*_ the number of cells below the node *k* and *N* the total number of cells, as described in Section 4.1. Hence, we would achieve significance at any α belonging to the set

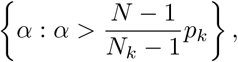

and we thus report the adjusted p-value as the infimum of this set, i.e. 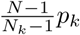.

### 4.3 Significance Analysis for Pre-Computed Clusters

We started by computing a tree-like hierarchy for any given set of clusters. Specifically, for each of *K* clusters, we computed the average expression for an informative subset of genes. These can either be the set of genes used in the clustering algorithm, if available, or they can be chosen as the *G* = 2, 500 genes with the largest deviance under a multinomial null model [12]. Because there are many cells per cluster, the data are no longer small counts for these centers, and we applied Euclidean distance to obtain a *K* × *K* distance matrix. This created a tree whose leaves are the cluster labels, and we followed a similar procedure as before.

In particular, we began at the root node and considered the two-way split dividing all cells into two clusters. As before, we applied the approximate GLM-PCA transformation, computed Euclidean distance using 30 latent factors, and computed the average silhouette value *s*. We then fit the parametric model to the data and performed parametric boostrap to generate counts for each cell under the null hypothesis. Because we rarely have access to the exact clustering algorithm used to produce the original clusters, we assigned the randomly generated cells to a cluster by first transforming them with the same GLM-PCA transformation applied to the observed data, computed a distance between each observed cell and each boostrapped cell, and, for each bootstrapped cell, used majority voting from the k-nearest neighbor observed cells. This provided cluster assignments for each bootstrapped sample, which in turn permitted us to compute a null distribution and proceed as in Section 4.1.

### 4.4 scRNA-seq gating plots

To visualize differences at proposed splits in hierarchical clustering, we introduced *scRNA-seq gating plots*, inspired by the dot plots used in flow cytometry. At any given node, we first identified differentially expressed genes between cells in the two clusters in question. We used the `findMarkers` function implemented by `scran` [26] and considered the set of genes with FDR *<* 0.05 and absolute log-fold-change greater than 0.5. For each cell, we then computed two quantities: the proportion of UMIs attributed to genes in this set that are more highly expressed in the first cluster, and the proportion of UMIs attributed to genes in this set that are more highly expressed in the second cluster. The scRNA-seq gating plots compare these two quantities for all cells, colored by cluster label, with density plots on the margins. If the two clusters in fact represent distinct populations, one should be able to visually see at least two clusters in this plot.

## Supplementary Methods

### 1 Simulated Study

To investigate over-clustering in current clustering workflows, we simulated scRNA-seq data from a single distribution to represent one cell population. In particular, we simulated 5,000 cells with 1,000 genes each, representing expressed genes. Previous exploratory analyses found the Poisson log-normal distribution to be appropriate for expressed genes [13], so counts for gene *g*, where *g* = 1, …, 1000, were drawn from a Poisson log-normal(*μ*_*g*_, 3) distribution. The parameters *μ*_*g*_ were set by sampling from a Normal(0, 2) distribution and then fixing those values for all cells.

These data were clustered using Seurat’s implementation of the Louvain algorithm [2]. We applied a standard workflow consisting of the following Seurat functions: `NormalizeData`, `ScaleData`, `RunPCA`, `RunUMAP`, `FindNeighbors`, and `FindClusters`. All functions were run using default parameters, except for `FindClusters`, which we ran 10 separate times with resolution parameters varying from 0.1 to 1, with intervals of 0.1. Note that the default resolution parameter is 0.8. We then used a popular clustering stability analysis, `clustree` [6], to visualize the results across these different parameter choices.

### 2 Parametric Model of Gene Expression

Define the *G × N* UMI counts matrix as ***Y***. Motivated by previous work [13], we assumed each gene *g* belongs to one of two latent states: unexpressed or expressed. If *J*_0_ is the set of indices corresponding to unexpressed genes and *J*_1_ is the set of indices corresponding to expressed genes, then

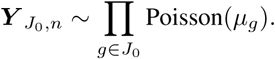

This implies that unexpressed genes are described by independent Poisson distributions with means *μ*_*g*_. The expressed genes were assumed to follow a Poisson log-MVN distribution:

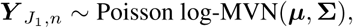

with mean vector ***μ*** of length |*J*_1_| and |*J*_1_| *×* |*J*_1_| covariance matrix **Σ**, for all cells *n* = 1, …, *N*. When considering all *G* genes, this forms a joint distribution for the cell population, which we refer to as ℱ.

### 3 Fitting the Parametric Model

Given observed ***Y***, we can fit *F* in a computationally efficient way using Method of Moments. First, we determined which genes belong to which latent state by testing for overdispersion. Specifically, we fit a Poisson GLM for each gene, then specified the variance as *μ*_*g*_ + *αμ*_*g*_ and tested the one-sided hypothesis *α* = 0 versus *α >* 0 at level 0.05. Genes that are overdispersed, which corresponds to rejecting this null hypothesis, were assumed to be expressed, and otherwise they were assumed to be unexpressed.

Next, we estimated the parameters of *F* using expressions derived from the moment equations. For each unexpressed gene *g*, we set

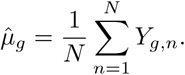

For each expressed gene *g*, we set

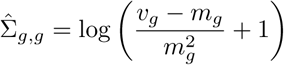

and

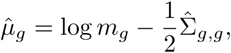

where *m*_*g*_, *v*_*g*_ are the sample mean and variance respectively for gene *g*. Then for any two genes *g*_1_, *g*_2_ such that *g*_1_ ≠ *g*_2_, we set

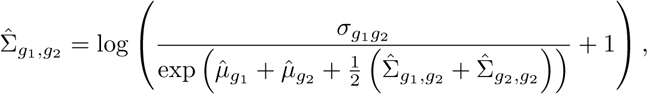

where *σ*_*g*1 *g*2_ is the sample covariance between genes *g*_1_ and *g*_2_.

Finally, because the resulting covariance matrix was not guaranteed to be positive definite, we approximated 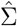 as the closest positive definite matrix under the infinity norm using the algorithm from [27], with the constraint that the variances, represented by the diagonal elements, are preserved.

### 4 Quantifying Clustering Quality with the Average Silhouette

Given two clusters, we quantified the clustering quality by computing the average silhouette [14], which is found as follows. If *C*_1_ and *C*_2_ denotes the indices for the cells belonging to clusters 1 and 2, respectively, then for each *i*, we compute

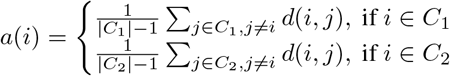

where *d*(*i, j*) is the distance between cells *i, j*. This represents the mean distance between cell *i* and all other cells in its cluster. We also computed

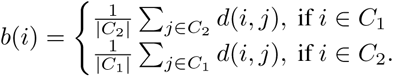

This now represents the mean distance between cell *i* and all cells in the other cluster. The silhouette of cell *i* is then written as

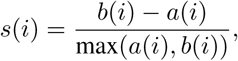

such that *s*(*i*) is close to 1 when *a*(*i*) ≪ *b*(*i*), indicating that cell *i* is clustered well; *s*(*i*) is close to *−*1 when *b*(*i*) ≪ *a*(*i*), indicating that cell *i* should belong to the other cluster; and *s*(*i*) is close to 0 when there is little difference between cell *i*’s distances to either cluster. We thus found the average silhouette for the entire clustering structure as

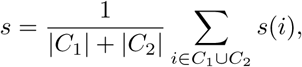

where values closer to 1 indicate higher quality clustering overall.

### 5 Benchmark Data

To construct data for benchmarking, we used 2,885 cells from the 293T cell line [16]. The one-population dataset consisted of these 2,885 cells with no changes. To construct the five-populations dataset, the first 20% of the cells were unaltered. Then, distinct permutations over a set of 100 genes, representing the 50 highest-expressing and 50 lowest-expressing genes, were applied to each successive 20% of the cells. The five-similar-populations dataset was created using this same process, except the permutation for the last 20% of the cells differs by only ten genes from the permutation used for the previous 20% of the cells.

When applying the Louvain and Leiden algorithms as implemented by Seurat, we ran these algorithms in two ways. First, we used the default resolution parameter of 0.8. Second, we used `clustree` to pick the optimal resolution parameter. This was done by varying the resolution parameter from 0.1 to 1 at intervals of 0.1, then choosing the largest parameter such that none of the clusters from the next resolution overlap by more than 25% of cells with two different clusters from the resulting output.

## Notes

### Competing Interest Statement

The authors have declared no competing interest.

